# Progressive heterogeneity of enlarged and irregularly shaped apicoplasts in *P. falciparum* persister blood stages after drug treatment

**DOI:** 10.1101/2024.01.03.574077

**Authors:** Chiara E. Micchelli, Caroline Percopo, Maria Traver, Joseph Brzostowski, Shuchi N. Amin, Sean T. Prigge, Juliana M. Sá, Thomas E. Wellems

**Author notes:** Thomas E. Wellems, **Email:**. **Author Contributions:** Designed research: CEM, JMS, TEW Performed research: CEM, CP, MT, JB, SNA Contributed new reagents or analytic tools: STP Analyzed data: CEM, JMS, TEW Wrote paper: CEM, TEW. **Competing Interest Statement:** We declare no competing interests. **Classification:** Biological, Health, and Medical Sciences – Cell Biology.

## Abstract

Morphological modifications and shifts in organelle relationships are hallmarks of dormancy in eukaryotic cells. Communications between altered mitochondria and nuclei are associated with metabolic quiescence of cancer cells that can survive chemotherapy. In plants, changes in the pathways between nuclei, mitochondria, and chloroplasts are associated with cold stress and bud dormancy. *Plasmodium falciparum* parasites, the deadliest agent of malaria in humans, contain a chloroplast-like organelle (apicoplast) derived from an ancient photosynthetic symbiont. Antimalarial treatments can fail because a small fraction of the blood stage parasites enter dormancy and recrudesce after drug exposure. Altered mitochondrial-nuclear interactions in these persisters have been described for *P. falciparum*, but interactions of the apicoplast remained to be characterized. In the present study, we examined the apicoplasts of persisters obtained after exposure to dihydroartemisinin (a first-line antimalarial drug) followed by sorbitol treatment, or after exposure to sorbitol treatment alone. As previously observed, the mitochondrion of persisters was consistently enlarged and in close association with the nucleus. In contrast, the apicoplast varied from compact and oblate, like those of active ring stage parasites, to enlarged and irregularly shaped. Enlarged apicoplasts became more prevalent later in dormancy, but regular size apicoplasts subsequently predominated in actively replicating recrudescent parasites. All three organelles, nucleus, mitochondrion, and apicoplast, became closer during dormancy. Understanding their relationships in erythrocytic-stage persisters may lead to new strategies to prevent recrudescences and protect the future of malaria chemotherapy.

**Significance Statement:** Dormancy of blood-stage malaria parasites (as persister forms) frequently undermines treatment and may facilitate the evolution of drug resistance. Here, we examine changes that occur in dormancy with two *P. falciparum* organelles relative to the nucleus: the mitochondrion and the plastid-like apicoplast. As previously reported, the mitochondrion of persisters is consistently enlarged, irregularly shaped, and shifted into close apposition with the nucleus. However, apicoplasts exhibit a greater variety of shapes, volumes, and relative positioning during dormancy: some persisters maintain a regular appearing apicoplast, while others show dramatically altered apicoplasts, reminiscent of the chloroplast swelling and degradation that occurs with death from reactive oxygen species in various plant cells. Improved understanding of these processes will support new approaches in antimalarial chemotherapy.

## Introduction

Malaria remains a major public health burden, particularly in sub-Saharan Africa where most morbidity and mortality from the disease are caused by *Plasmodium falciparum*. Artemisinin and its derivatives (collectively termed ART herein) are first-line antimalarial drugs and serve as the foundation of artemisinin-based combination therapies (ACTs) for uncomplicated malaria (1, 2).

Although ART monotherapy rapidly clears the symptoms and parasitemia of malaria, combination treatment with a long-acting partner drug is necessary to prevent parasite recrudescences that often follow 3-, 5-, 7-, and even 10-day regimens of ART alone (3–7). Pharmacokinetic-pharmacodynamic modeling based on clinical outcomes after artesunate (AS) monotherapy suggested that a small number of parasites dodge the lethal effects of ART by entering a state of dormancy (8). Indeed, inactive dormant forms of *P. falciparum* had been proposed from studies of parasite cultures subjected to repetitive selections with sorbitol or treatments with pyrimethamine (PYR) or mefloquine (MEF) (9, 10). Subsequent studies treating parasites in vitro with either artemisinin or dihydroartemisinin (DHA) showed that a subpopulation of ring-stage parasites can also survive by entering a metabolically quiescent, growth arrested state (11–13). These dormant persister forms occur in human patients after single doses of AS (14). *Plasmodium* dormancy has also been reported after exposure to other antimalarial drugs including chloroquine (CQ) (15, 16) as well as atovaquone alone or in combination with proguanil (17).

Morphological modifications and shifts in organelle relationships are hallmarks of cellular stress and dormancy in eukaryotic cells. In plants, condensation of chromatin and increased contact between nuclei, mitochondria, and chloroplasts are associated with bud dormancy and cold stress (18–20).

Chloroplasts are especially sensitive to abiotic stress and reactive oxygen species, which can alter inter-organelle communication and cause chloroplast transformations, including swelling and degradation (21, 22). In mammals, mitochondrial changes are common markers of cellular stress. So called ‘giant mitochondria’ are found in the liver parenchymal cells of patients with non-alcoholic fatty liver disease, and mitochondrial-nuclear associations in cancer cells may alter retrograde signaling and thereby facilitate survival (23, 24). In cancer cells treated with cisplatin, nuclear pore architecture has also been shown to change with quiescence (25). Cell subpopulations in a stress-like, quiescent state occur in early stages of tumorigenesis with several types of cancers. Innate groups of these stress-like cells can seed new tumors and cause disease relapses after chemotherapy, independent of mutations (26). Dormancy in yeast is accompanied by changes including extensive reorganization and compaction in the cytoplasm (27).

Previous work from our group demonstrated dramatic changes in volume and localization of mitochondria in dormant blood stages of *P. falciparum*. The mitochondrion in these stages enlarges and forms a close association with the nucleus, potentially facilitating organelle contact sites and retrograde signaling pathways common in cellular stress (28). Another organelle that has been implicated in *P. falciparum* dormancy is the apicoplast, a plastid organelle derived from the endosymbiosis of red algae (29–31). Here we use super-resolution Airyscan microscopy (ASM) to examine the status of the apicoplast in *P. falciparum* blood stage persisters and active ring stages. We compare the apicoplast and mitochondrion volumes and the spatial relationships of these organelles with respect to each other and to the nucleus in dormancy.

## Results

### Isolation of persisters and active ring stages using fluorescence-activated cell sorting

To study organelle morphology in dormant blood-stage parasites, we chose the NF54-SFG parasite line that expresses a fluorescent Super Folder Green label in the apicoplast (NF54-SFG) (32). The dormancy and recrudescence curves of this line (**Figure S1**) were obtained by the method of Breglio *et al.* (33), in which synchronized ring stages were treated with DHA for 6 h on day 0, followed by 5% sorbitol selection at 24, 48, and 72 h (days 1, 2, and 3) to remove the trophozoites and schizonts of any remaining actively replicating parasites by osmotic lysis (34). The parasitemia of active forms (microscopically identified ring-, trophozoite-, and schizont-stage parasites) rapidly decreased after DHA treatment on day 0, and by the third sorbitol treatment only persisters and dying or dead (pyknotic) parasites were observed. Persisters were distinguished by rounded, compact chromatin with condensed cytoplasm visible in Giemsa-stained thin blood smears, as previously described (35). Active forms became visible between days 11 and 13, and the parasitemia rose to ≥ 2% between day 16 and 17.

To isolate and enrich NF54-SFG parasite-infected erythrocytes for the examination of apicoplast morphology, active ring stages and persisters were stained with Hoechst 333432 (HO) and MitoTracker Deep Red (MT) and collected by fluorescence-activated cell sorting (FACS) in a two-step enrichment protocol (see Methods). Persisters were obtained by treating synchronized ring-stage parasites with 700 nM DHA/0.1% DMSO in culture medium for 6 h on day 0, followed by 5% sorbitol on day 1 to eliminate the possible presence of parasites that were not dormant and had progressed from active ring forms to more mature stages. For control observations, synchronized parasites were treated with 0.1% DMSO (the drug vehicle) in culture medium on day 0 and mock-selected with incomplete culture medium (no sorbitol, see Methods section) on day 1; in this report, these are referred to as “vehicle-treated control parasites” or “controls” (**Figure S2**).

Cells were collected from culture on day 2, stained with HO and MT, and separated using the FACS gating strategy shown in **Figure 1A**. In the first of a two-step sorting process, the yield sort enriched samples for parasitized erythrocytes based on HO signal, thereby removing a significant proportion of uninfected red cells. In the second (purity) sort, infected erythrocytes were further separated from uninfected cells based on both HO and MT fluorescence. HO+MT+ cells (shown in purple) were gated into four subpopulations based on their different levels of MT fluorescence (P1, P2, P3, and P4).

**Figure 1.**
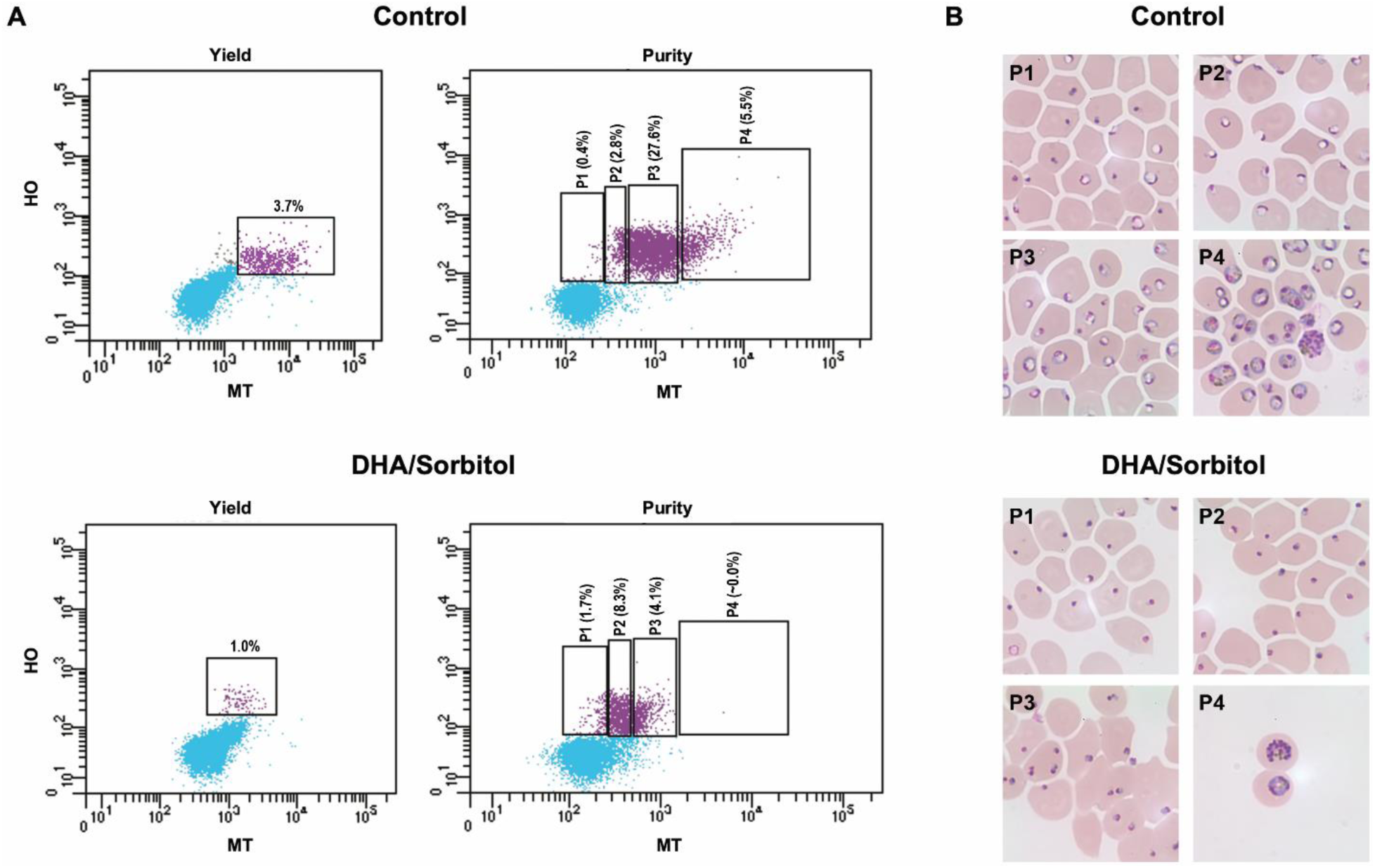
FACS gating strategy and Giemsa-stained cytospin images of control- and DHA/sorbitol-treated NF54-SFG parasites. Synchronized 6–9 h rings were exposed to the drug vehicle for 6 h on day 0 and mock-selected with iRPMI on day 1 (Control), or they were treated with 700 nM DHA for 6 h on day 0 and selected with 5% sorbitol on day 1 (DHA/Sorbitol). The samples were harvested on day 2 (t= 50 h), stained with Hoechst 33342 (HO) plus MitoTracker Deep Red FM (MT). Parasitized erythrocytes were collected and imaged after two sequential steps of purification by the yield and 4-way purity modes of a BD Aria™ Fusion Flow Cytometer. (A) Panels labeled ‘Yield’ show the parasitemias of culture samples and gates employed for the first step of high-throughput yield sorting. Panels labeled ‘Purity’ show the distributions of cells obtained after yield sorting and delineate the P1 – P4 gates used to collect the purity-sorted populations. Uninfected red cells are represented in blue and HO+/MT+ cells in purple. The vehicle-, iRPMI mock-treated population containing 3.7% HO+ cells was yield-purified, providing an enriched distribution of 0.4%, 2.8%, 27.6%, and 5.5% HO+ cells in the P1 – P4 windows for purity-collected controls. Similarly, the population of 1% HO+ cells from the DHA/sorbitol-treated culture was yield-enriched to provide 1.7%, 8.3%, 4.1%, and very rare cells (0.0% reported) in the P1 – P4 windows for purified DHA/sorbitol-treated cells. (B) Images of Giemsa-stained parasites from cytospin preparations of P1–P4 subpopulations. Late rings and trophozoites predominate in the P3 subpopulation of control-treated samples; smaller percentages of late-stage schizonts and early rings are explained by loss of synchrony from experimental manipulations or naturally arising cell-to-cell variability (96). In DHA/sorbitol-treated samples, persisters were most frequent in the P2 window. Although FACS detected almost no events in the DHA/Sorbitol P4 window, our searches identified rare mature forms in highly concentrated cytospin preparations (two parasitized cells shown).

Cells with the lowest MT fluorescence fell into the P1 gate, and cells with the highest MT fluorescence were located in the P4 gate. In these separations, the cells of vehicle-treated control parasites were distributed across all levels of MT positivity, with late rings and trophozoites predominating in the P3 subpopulation. This was expected as, in our laboratory cultures, the blood-stage cycle of NF54-SFG parasites was approximately 40 h, consistent with the 39 h cycle time found in human volunteers (36). Thus, after the time to harvest (50 h) plus another ∼6 h to complete the FACS and cytospin preparations, the parasites imaged in the upper panels were well into their second 40 h cycle, at 22 – 25 h development (**Figure 1B; Figure S2**).

Images of Giemsa-stained cells collected by cytospin from each FACS-isolated subpopulation are shown in **Figure 1B**. The P1 subpopulation of vehicle-treated control cells consisted of uninfected erythrocytes, some erythrocytes infected with rings, and erythrocytes infected with persisters including pyknotic-like forms. The P2 subpopulation consisted of erythrocytes infected with single early rings, while the P3 subpopulation included erythrocytes infected with single or multiple rings, late rings, and early trophozoites. P4, the subpopulation with the brightest MT signal, contained cells infected with schizonts, trophozoites, and multiple parasites.

On day 2, the subpopulation compositions of the DHA/sorbitol-treated parasites differed markedly from that of the controls (compare **Figure 1B**, Control vs. DHA/Sorbitol panels). Greater numbers of HO+MT+ cells were obtained from the controls than from the DHA/sorbitol-treated cultures. Cells in the DHA/sorbitol P1 subpopulation consisted mostly of pyknotic-like parasites. The P2 and P3 subpopulations contained erythrocytes infected with persisters. Erythrocytes in P3 were sometimes infected with more than one persister, but these multiple infections were typically not observed in the P2 cells. Finally, the P4 subpopulation contained sparse numbers of cells, making their collection and observation difficult. The few P4 cells that could be collected and examined microscopically contained parasites in advanced stages of development (trophozoites and schizonts).

### Altered morphology and localization of apicoplasts, as well as mitochondria, in parasites after DHA/sorbitol exposure

The organelles of parasites isolated on day 2 from the P2 and P3 subpopulations of DHA/sorbitol- and vehicle-treated control cultures were imaged using ASM. Organelle ‘surfaces’ were processed in Imaris based on 3-D reconstructions from Airyscan-processed images, which also enabled spatial and volumetric analysis. **Figure 2A** displays representative surface renderings of the fluorescence volumes of the mitochondrion (red), nucleus (blue), and apicoplast (green). **Figures 2B and 2C** show individual organelle volumes and distances, respectively, along with their corresponding medians and interquartile ranges (IQRs).

**Figure 2.**
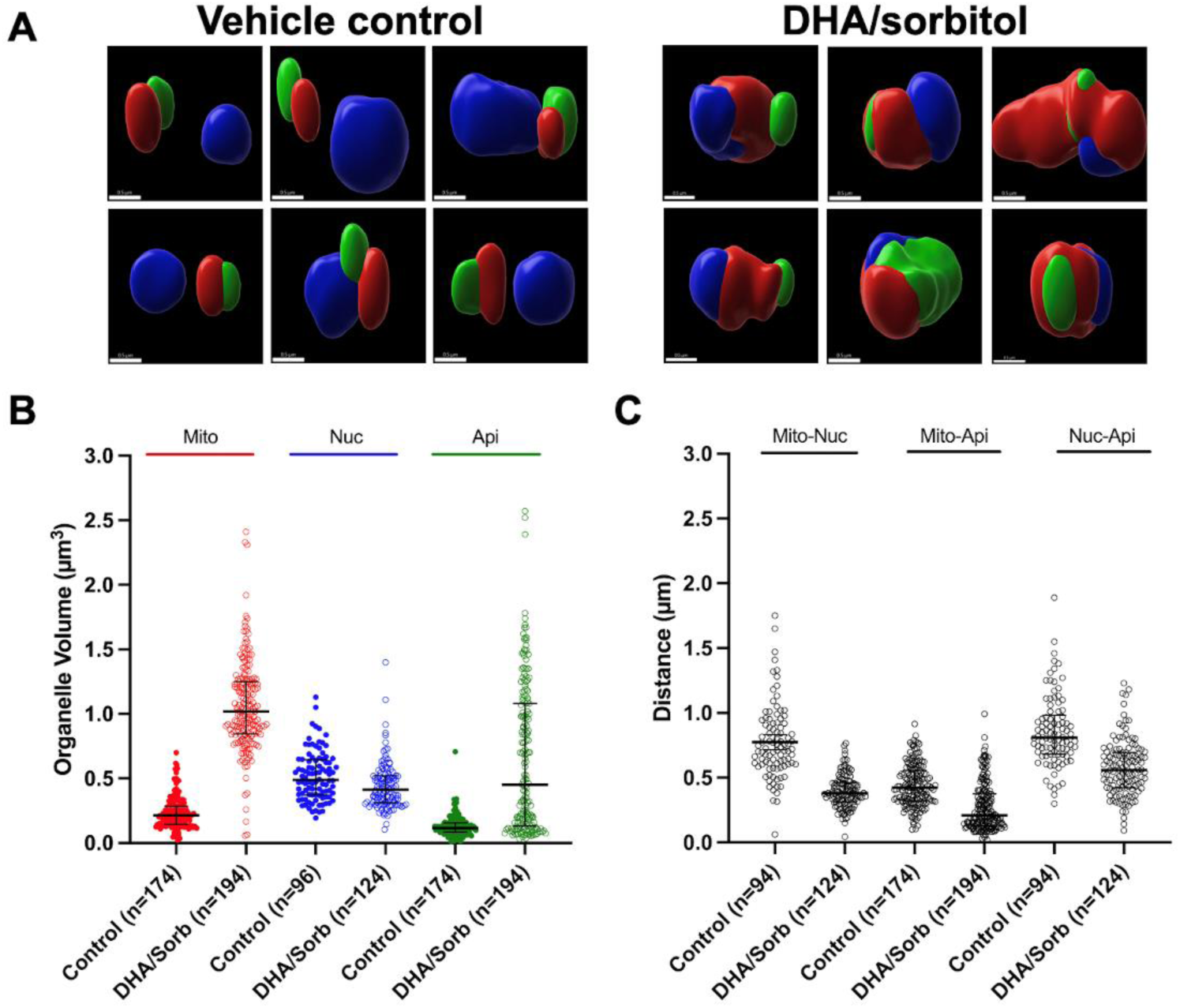
Organelle images and measurements from vehicle-treated control and DHA/sorbitol-treated parasites at day 2. The vehicle-treated control and DHA/sorbitol-treated parasites were isolated as described in Figure 1 and imaged using Airyscan microscopy. (**A**) Airyscan-processed fluorescence images of organelles in DHA/sorbitol-treated persisters show dramatic changes of morphology and association compared to the organelles of vehicle control parasites. Apicoplasts, mitochondria, and nuclei are represented in green, red, and blue, respectively. Surfaces were created using Imaris. Scale bar= 0.5 µm. (**B**) Scatterplots, medians, and interquartile ranges (IQRs) of apicoplast (Api), mitochondrial (Mito), and nuclear (Nuc) volumes in vehicle-treated active ring stages and DHA/sorbitol-treated persisters. Each point represents one organelle volume. Mann-Whitney significance results from comparisons of these volumes in vehicle control-*vs.* DHA/sorbitol-treated parasites: Mito P<0.0001, Nuc P<0.0009, Api P<0.0001 (**Dataset S1**). (**C**) Scatterplots, medians, and IQRs of the inter-organelle distances in vehicle-treated control and DHA/sorbitol-treated parasites. Centers of mass of the organelles were used to calculate the distances between the mitochondria and nuclei (Mito-Nuc), mitochondria and apicoplasts (Mito-Api), and nuclei and apicoplasts (Nuc-Api). Each point represents one distance measurement. Mann-Whitney significance test results from comparisons of these distance in vehicle control-*vs.* DHA/sorbitol-treated parasites: Mito-Nuc P<0.0001, Mito-Api P<0.0001, Nuc-Api P<0.0001 (**Dataset S1**).

In ring stages from the vehicle-treated control cultures, apicoplasts had a median volume of 0.12 µm^3^ (IQR 0.08 to 0.16 µm^3^) and were oblate in shape (**Figure 2A**, Vehicle control; **Figure 2B, Movie S1**; **Dataset S1**). The apicoplast in individual parasites was located immediately adjacent to the mitochondrion, consistent with the electron microscopy findings of Hopkins *et al.* (37). The median distance between these organelles, as measured from the center of mass of each organelle, was 0.42 µm (0.32-0.56 µm) (**Figure 2C, Dataset S1**). The apicoplast and mitochondrion were separated from the nucleus by a median distance of 0.81 µm (0.68-0.98 µm) and 0.72 µm (0.60-0.92 µm), respectively (**Figure 2C, Dataset S1**). In some instances, the mitochondrion and apicoplast were closer to the nucleus but their fluorescence volumes still occupied discrete locations (**Movie S2**).

The mitochondria of DHA/sorbitol-treated parasites on day 2 were consistently enlarged, in agreement with previous observations (28). **Figure 2A** shows the mitochondria of control- and DHA/sorbitol-treated parasites in red, two days post-DHA. Whereas the mitochondria of the active ring-stage parasites were oblate with a median volume of 0.21 µm^3^ (0.14-0.29 µm^3^), the mitochondria of the persisters were irregularly shaped with a median volume of 1.0 µm^3^ (0.84-1.3 µm^3^; P<0.001, Mann-Whitney test) (**Figure 2B, Dataset S1**). As previously observed, the mitochondrion was more closely associated with the nucleus in persisters, as shown by the decrease in the median distance from 0.72 µm (0.60-0.92 µm) in rings to 0.38 µm (0.31-0.46 µm) in persisters (P<0.0001, Mann-Whitney test) (**Figure 2C, Dataset S1**).

The morphology of the apicoplast on day 2 varied more than that of the mitochondrion in persisters following DHA/sorbitol treatment. A proportion of these persisters had a compact, oblate apicoplast like that observed in active ring stages, whereas many others contained large and irregularly shaped apicoplasts (**Figure 2A and B**). In all persisters, the apicoplasts were close to the mitochondria. In persisters with oblate apicoplasts, the apicoplast sometimes appeared to be enwrapped by the mitochondrion (**Movie S3**).

The Airyscan-processed images of parasites with large apicoplasts displayed overlapping apicoplast, mitochondrial, and nuclear fluorescence in the 2D and 3D projections, as well as in individual z-slices (**Movie S4**). Furthermore, the fluorescence of large apicoplasts was often expanded with less intensity. For the apicoplasts of all sizes, the median volume in the persisters was increased to 0.42 µm^3^ (0.13-1.1 µm^3^; P<0.0001, Mann-Whitney test), and the median mitochondrion-apicoplast distance was 0.21 µm (0.13-0.37 µm; P<0.0001, Mann-Whitney test), suggesting more extensive association than in the active ring stage parasites (**Figure 2C, Dataset S1**). Although the swollen mitochondrion was typically larger than the apicoplast in persisters, this was sometimes not the case in persisters containing particularly enlarged apicoplasts.

**Figure 3A–C** shows organelle surface renderings as well as organelle volumes and inter-organelle distances on days 5, 11, and 17 in persisters that received one DHA treatment on day 0 and three sorbitol treatments on days 1, 2, and 3. As with the apicoplasts of day 2 parasites after one DHA treatment and one sorbitol treatment, marked morphological diversity was evident among the apicoplasts in the day 5 and day 11 persisters. On day 5, the apicoplast shape and volume were often oblate and < 0.5 µm^3^, however abundant swollen forms were also observed with apicoplast volumes >0.5 µm^3^. Taken together, the median apicoplast volume of the DHA/sorbitol-treated day 5 forms was 0.65 µm^3^ (0.12-1.1 µm^3^), significantly higher than that of vehicle-treated control parasites on day 2 (P<0.0001, Kruskal-Wallis test). By day 11, there was an even greater departure from the apicoplast morphology characteristic of active rings (P<0.0001, Kruskal-Wallis test). Although some apicoplasts in the day 11 persisters continued to show regular oblate volumes, the majority of apicoplasts were enlarged and in some cases were larger than the mitochondria. Many of the persisters exhibited fluorescence overlap, especially from apicoplast and mitochondrial markers. These observations coincided with the findings of greatly increased median apicoplast volume (1.2 µm^3^ [0.91-1.5 µm^3^]; P<0.0001, Kruskal-Wallis test) and reduced distance between the mitochondrion and apicoplast (0.11 µm [0.09-0.14 µm]; P<0.0001, Kruskal-Wallis test), as well as between the apicoplast and nucleus (0.45 µm [0.33-0.55 µm]; P<0.0001, Kruskal-Wallis test; **Figures 3B and C, Dataset S1)**.

**Figure 3.**
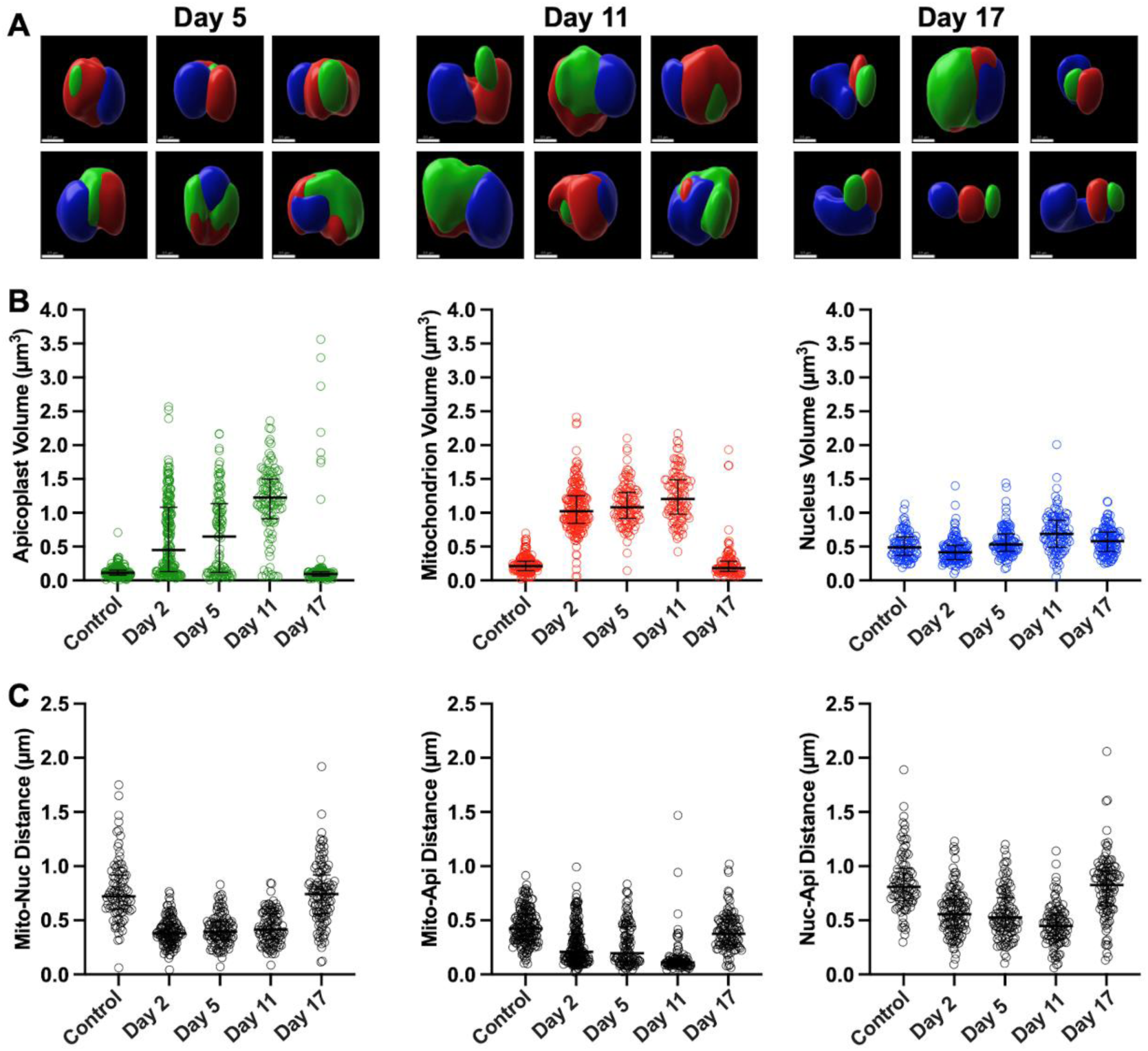
ASM images, organelle volumes, and inter-organelle distances of DHA/sorbitol-treated parasites on days 5, 11, and 17. Synchronized ring-stage parasites were treated with 700 nM DHA on day 0, followed by three daily 5% sorbitol treatments on days 1, 2, and 3. ASM analysis was performed on infected erythrocytes from the P2 and P3 subpopulations isolated by FACS on days 5, 11, and 17. (**A**) Representative images from parasites on days 5 and 11 show predominantly persister forms, whereas the images from parasites on day 17 almost exclusively show active ring stages. Scale bar= 0.5 µm. (**B and C**) Scatterplots with medians and IQRs of the organelle volumes and inter-organelle distances in DHA/sorbitol-treated parasites on days 5, 11, and 17. Results from vehicle-treated control and DHA/sorbitol-treated parasites on day 2 are included for reference. Kruskal-Wallis significance test results from comparisons of the volumes in vehicle control-*vs.* DHA/sorbitol-treated parasites: Apicoplast Day 2 P<0.0001, Day 5 P<0.0001, Day 11 P<0.0001, Day 17 P>0.9999; Mitochondrion Day 2 P<0.0001, Day 5 P<0.0001, Day 11 P<0.0001, Day 17 P>0.9999; Nucleus Day 2 P=0.0084, Day 5 P=0.3639, Day 11 P<0.0001, Day 17 P=0.0491. Kruskal-Wallis significance test results from the distance comparisons: Mito-Nuc Day 2 P<0.0001, Day 5 P<0.0001, Day 11 P<0.0001, Day 17 P>0.9999; Mito-Api Day 2 P<0.0001, Day 5 P<0.0001, Day 11 P<0.0001, Day 17 P>0.9999; Nuc-Api Day 2 P<0.0001, Day 5 P<0.0001, Day 11 P<0.0001, Day 17 P>0.9999 (**Dataset S1**).

**Figure 3B** shows that the mitochondria of persisters were more uniformly enlarged than the apicoplasts at days 2, 5, and 11. The median mitochondrial volume was 1.08 µm^3^ on day 5 (0.92-1.3 µm^3^; P<0.0001, Kruskal-Wallis test) and 1.2 µm^3^ on day 11 (0.98-1.5 µm^3^) (P<0.0001, Kruskal-Wallis test).

Along with these changes of volume, there was a reduction in the median distance between the apicoplast and the nucleus (**Figure 3C; Dataset S1**). The nuclear volumes displayed comparatively small changes in persisters. Two days post-DHA, the median nuclear volume was 0.41 µm^3^ (0.31-0.52 µm^3^), less than that of ring stages (0.49 µm^3^ [0.37-0.64 µm^3^]; P=0.0009, Mann-Whitney test), but it increased on days 5 and 11 to 0.53 µm^3^ (0.43-0.69 µm^3^; P=0.3639, Kruskal-Wallis test) and 0.69 µm^3^ (0.49-0.89 µm^3^; P<0.0001, Kruskal-Wallis test), respectively (**Figure 3B, Dataset S1**).

At recrudescence, on day 17 post-DHA, relatively few persisters remained, and active ring stages were the most prevalent in the P2 and P3 FACS subpopulations. ASM analysis and Imaris-generated surfaces showed that all organelles in the day 17 ring stages were morphologically similar to those of ring stages from the vehicle-treated control cultures on day 2 (**Figure 3A**, Day 17). The median apicoplast, mitochondrial, and nuclear volumes of the recrudescent ring stages were 0.09 µm^3^ (0.06-0.13 µm^3^), 0.18 µm^3^ (0.14-0.28 µm^3^), and 0.58 µm^3^ (0.43-0.72 µm^3^) respectively (**Figure 3B, Dataset S1**). These median apicoplast and mitochondrial volumes were not significantly different from those of vehicle-treated control parasites (P>0.999 in both cases, Kruskal-Wallis test), while the p-value for a somewhat larger median nuclear volume of recrudescent *vs.* control ring stages (0.58 µm^3^ vs 0.48 µm^3^) was marginal (P=0.0491, Kruskal-Wallis test). This larger nuclear volume could be due to asynchronicity of ring stages upon recrudescence, whereas rings in the controls were tightly synchronized prior to collection. The spatial relationships between organelles in the recrudescent ring stages also returned to pre-DHA/sorbitol treatment values (**Figure 3C, Dataset S1**). The few persisters that remained on day 17 displayed characteristically enlarged apicoplasts and mitochondria with overlapping fluorescence signals (**Figure 3A**).

### Apicoplast volume distributions in persisters over time

To comparatively evaluate the apicoplast changes over time, we grouped the DHA/sorbitol-treated persisters into five categories based on apicoplast volume: <0.5 µm^3^, 0.5-1.0 µm^3^, 1.0-1.5 µm^3^, 1.5-2.0 µm^3^, and >2.0 µm^3^. The percentages of the parasites containing apicoplasts in these categories were determined from the populations of three biological replicate experiments as well as vehicle-treated controls. **Figure 4** shows a heat map of apicoplast prevalence in these size ranges on days 2, 5, 11, and 17 of the recrudescence experiments plotted in **Figure S1**.

**Figure 4.**
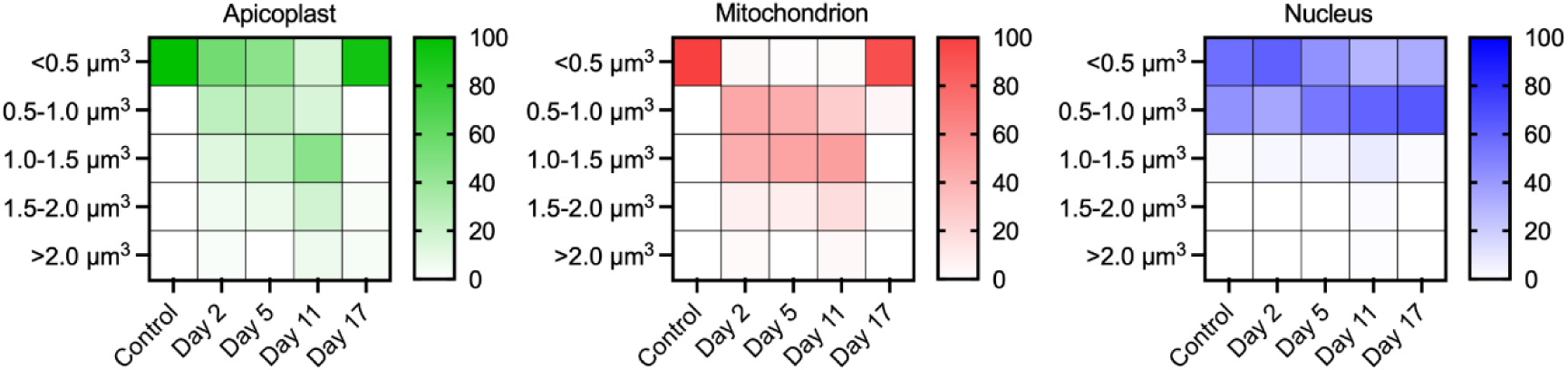
“Heat maps” showing changes of the organelle volume distributions in DHA/sorbitol-treated persisters over time. The grid represents mean apicoplast, mitochondrion, and nuclear volumes of vehicle-treated control and DHA/sorbitol-treated parasites on days 2, 5, 11, and 17 obtained from three biological replicates. The organelles were categorized into five groups based on volume: <0.5 µm^3^, 0.5-1.0 µm^3^, 1.0-1.5 µm^3^, 1.5-2.0 µm^3^, and >2.0 µm^3^.

In the control cells on day 2, only regular <0.5 µm^3^ volume apicoplasts were observed in active ring stages. The mean prevalence of <0.5 µm^3^ apicoplasts in DHA/sorbitol-treated persisters on day 2 was 55.1 ± 9.38% (SEM), with progressively lower prevalences in larger volume categories (**Figure 4, Figure S3, Dataset S1**). On day 5 after DHA/sorbitol, the <0.5 µm^3^ group continued to be most prevalent (45.4 ± 15.1%, Mean ± SEM) but with a trend to increased overall counts of larger apicoplast volumes. On day 11, the swollen apicoplasts predominated, with mean prevalences of 45.6 ± 8.9% and 18.0 ± 5.8% in the 1.0-1.5 µm^3^ and 1.5-2.0 µm^3^ groups, respectively. The mean prevalence of >2.0 µm^3^ volume apicoplasts was also highest on day 11 (6.7% ± 3.3%). Even though these large volumes were most abundant, we note that the <0.5 µm^3^ apicoplasts remained prevalent (14.9 ± 5.7%). As described above, nearly all apicoplasts in the FACS-sorted samples on day 17 were <0.5 µm^3^ (93.0% ± 7.0%; **Figure S3, Dataset S1**) and exhibited ring-stage morphology, consistent with the presence of abundant actively replicating parasites in the recrudescent population.

Corresponding analyses were performed on the mitochondrial and nuclear volumes (**Figures 4 and S3, Dataset S1**). Like the apicoplast volumes, the mitochondrial volumes in vehicle-treated control parasites were uniformly <0.5 µm^3^. In the DHA/sorbitol treated persisters, however, most mitochondria became enlarged: on days 2, 5, and 11, 45.2 ± 11.1%, 42.7 ± 11.7%, and 25.8 ± 9.7% of the calculated mitochondrial volumes were 0.5-1.0 µm^3^, and 42.4 ± 8.3%, 47.5 ± 7.3%, and 50.4 ± 1.49% were 1.0-1.5 µm^3^, respectively. By day 17, mitochondria with <0.5 µm^3^ volumes predominated again (92.7 ± 5.63% <0.5 µm^3^), consistent with the recrudescence of actively replicating parasites. Nuclear volumes showed less variation between the vehicle-treated control and the DHA/sorbitol-treated parasites across the different days after drug treatment.

### Dormancy, recrudescence, and changes in apicoplast morphology observed after Sorbitol treatment alone

In addition to its use as a synchronization agent for *P. falciparum* cultures, sorbitol treatment alone may be used to study recrudescence from dormant blood-stage parasites in culture (9). To explore this phenomenon further, we subjected ring-stage parasites to 5% sorbitol every 24 h for 4 days.

Recrudescences after treatment with this protocol (4DS) arose several days sooner than after treatment with DHA/sorbitol (**Figure 5A**). Active ring- and trophozoite-stage parasites were seen 5-7 days after the first sorbitol treatment (1-3 days after the final sorbitol treatment) and parasitemias reached ≥2% on days 10-13.

**Figure 5.**
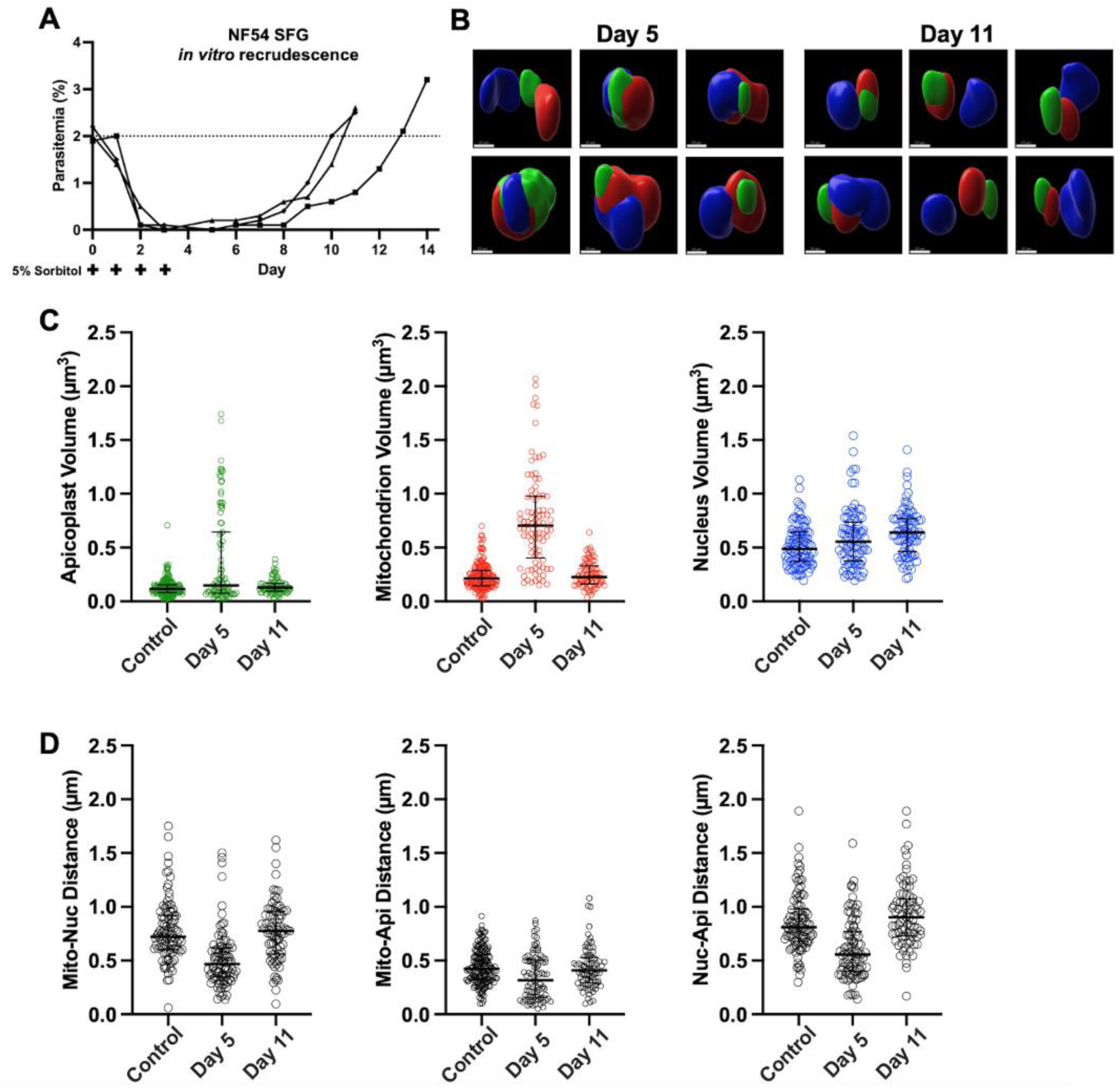
Recrudescence curves, ASM images, and organelle measurements of parasites after four daily sorbitol treatments. Synchronized ring-stage parasites were subjected to 5% sorbitol on day 0 for 10 min, followed by three additional sorbitol treatments for 30 min on days 1, 2, and 3. (**A**) Recrudescence curves of NF54-SFG active-form blood stages subjected to the four 5% sorbitol treatments. Parasitemias were monitored daily by Giemsa-stained thin blood smears until ≥ 2% recrudescence. Each line represents one biological replicate. (**B**) Sorbitol-treated cultures were stained with HO and MT, collected by FACS, and subjected to ASM analysis. Organelle surfaces and approximations on day 5 were similar to those of DHA/sorbitol-treated parasites on day 2. By day 11, most parasites contained apicoplasts, mitochondrion, and nuclei characteristic of active ring stages. Scale bar= 0.5 µm. (**C and D**) Scatterplots with medians and IQRs of organelle volumes and inter-organelle distances in the sorbitol-treated parasites. Kruskal-Wallis significance test results from comparisons of the volumes in vehicle control- *vs.* sorbitol treated-parasites: Apicoplast Day 5 P=0.0015, Day 11 P=0.3540; Mitochondrion Day 5 P<0.0001, Day 11 P=0.4935; Nucleus Day 5 P=0.01525, Day 11 P=0.0003. Kruskal-Wallis significance test results from the distance comparisons: Mito-Nuc Day 5 P<0.0001, Day 11 P>0.9999; Mito-Api Day 5 P=0.0001, Day 11 P=0.8707; Nuc-Api Day 5 P<0.0001, Day 11 P=0.2543 (**Dataset S1**).

**Figure 5B** shows representative surface renderings from ASM images of the 4DS-treated parasites on days 5 and 11, and **Figure 5C and D** compares the corresponding organelle volumes and distances of vehicle-treated control *vs.* 4DS-treated parasites. On day 5, the 4DS-treated persisters exhibited organelle morphologies similar to those of DHA/sorbitol-treated persisters on day 2, with marked decreases in the median mitochondrial-nuclear, mitochondrial-apicoplast, and nuclear-apicoplast distances relative to those of active ring-stage parasites (**Figure 5D, Dataset S1**). Counts of the 4DS-treated persister organelle volumes reflected their diversity in size at day 5: the apicoplasts in 63 of 89 persisters were regularly oblate with volumes < 0.5 µm^3^, whereas apicoplasts in the remaining 26 of 89 contained large, swollen apicoplasts having volumes >0.5 µm^3^. Taking these apicoplast measures together, the median volume was 0.14 µm^3^ (0.08-0.63 µm^3^), significantly higher than the control (P=0.0015, Kruskal-Wallis test). The mitochondrial volumes of the 4DS-treated persisters were highly diverse, with a median mitochondrial volume of 0.71 µm^3^ (0.40-0.98 µm^3^), greater than median mitochondrial volume of active rings (P<0.0001, Kruskal-Wallis test), but less than that of DHA/sorbitol-treated parasites on days 2 and 5. The median nuclear volume was not significantly different from that of the control (0.56 µm^3^ [0.38-0.74 µm^3^]; P=0.1525, Kruskal-Wallis test) (**Dataset S1)**.

By day 11, many of the 4DS-treated parasites were actively replicating, consistent with recrudescence from persisters. Ring stages and remaining persisters in the P2-P3 gating windows were collected and imaged. The apicoplasts and mitochondria in these collections exhibited oblate morphologies indistinguishable from those of untreated rings on day 2. The median apicoplast volume was 0.13 µm^3^ (0.09-0.17 µm^3^), the median mitochondrial volume was 0.23 µm^3^ (0.16-0.33 µm^3^), and the median mitochondrial-apicoplast distance was 0.41 µm (0.29-0.53 µm) (**Figure 5C and D, Dataset S1**). The median nuclear volume on day 11 was larger than that of rings on day 2 (0.64 µm^3^ [0.46-0.78 µm^3^]; P=0.0003, Kruskal-Wallis test), but the median distances from nuclei to mitochondria (0.78 µm [0.56-0.96 µm]) and nuclei to the apicoplasts (0.90 µm [0.73-1.1 µm]) were not significantly different from those of the control (P>0.9999 and P=0.2543, respectively; Kruskal-Wallis test).

## Discussion

In this study we examined the morphology of the apicoplast organelle in *P. falciparum* blood-stage persisters obtained after exposure to DHA and sorbitol (DHA/sorbitol) or to sorbitol alone. In ASM images, the apicoplasts of many persisters appeared small and oblate, similar in size to the apicoplasts of active ring-stage parasites, while the apicoplasts of others were swollen and irregular in shape. The apicoplasts of all persisters localized near the enlarged and corrugated mitochondria previously described by Connelly *et al.* (28). After DHA/sorbitol exposure, the small oblate apicoplasts were observed more frequently than swollen forms on days 2 and 5, then the swollen forms predominated on day 11. On day 17, after exit from dormancy and recrudescence, a majority of actively replicating parasites were observed again, with small and oblate apicoplasts in the ring stages. The appearance and median volume of mitochondria were also normal in the active ring stages on day 17. In persisters obtained by daily sorbitol treatments for four consecutive days, the organelle changes were similar to those seen after the use of DHA/sorbitol, but earlier recrudescences occurred at days 10–13.

The apicoplast is a non-photosynthetic organelle with important roles in the production of isoprenoids, fatty acids, iron-sulfur clusters, heme, and coenzyme A (38–40). It has also been tied to dormancy in blood stages of *P. falciparum.* The expression of several apicoplast-localized enzymes involved in fatty acid synthesis (FASII) and pyruvate metabolism was shown to be upregulated two days post-DHA treatment, despite generalized reductions in transcription and translation during dormancy (30, 41). Duvalsaint and Kyle (31) reported additional evidence for involvement of apicoplast lipid pathways from experiments showing that recrudescence after dormancy did not occur with parasites that had lost functional apicoplasts. Only upon addition of isopentenyl pyrophosphate (IPP), the most important apicoplast product, did persisters recover after antibiotic exposure. The same study found that treating persisters with gibberellic acid, a plant hormone that promotes emergence from seed dormancy and germination, caused early recrudescence, while treatment with an herbicide, fluridone, blocked recrudescence.

Reflecting on their evolutionary past, what additional features do *Plasmodium* parasites have in common with plants beyond the influence of plant hormones on dormancy? The apicoplasts of *Plasmodium* and *Toxoplasma* and chloroplasts of plants likely originated from the engulfment of cyanobacteria in primordial endosymbiotic events (42). Chloroplasts, which are major sources of reactive oxygen species (ROS), are especially sensitive to abiotic stressors and act as barometers of cellular health in plants. Changes in chloroplast morphology and organelle associations often occur during stress. For instance, stromules, stroma-filled extensions of chloroplasts, can form to facilitate retrograde signaling to nuclei in *Arabidopsis thaliana* upon exposure to ROS (43, 44). Chloroplasts, mitochondria, and nuclei have been observed to move into close proximity to each other in cold-stressed leaf mesophyll cells of Arctic plants (18). Our results showing closer proximity between apicoplasts, mitochondria, and nuclei in *P. falciparum* persisters are analogous to the findings of organellar repositioning in dormant plant cells, in which such changes may facilitate pro-survival responses to stress (18, 23, 43, 44).

The enlarged apicoplasts in *P. falciparum* persisters after DHA/sorbitol or sorbitol exposure are reminiscent of the swollen chloroplasts of plant cells exposed to heat or cold stress, high intensity light, high salinity, or osmotic imbalances (45–48). Swollen chloroplasts of *A. thaliana* are less efficient in carrying out photosynthesis than their smaller, undamaged counterparts (49), and are ultimately degraded in autophagy (ATG)-dependent chlorophagy mediated by ATG8 (22, 48). Interestingly, ATG8 localizes to the outer membrane of the apicoplast in *Plasmodium* and contributes to apicoplast biogenesis in blood and liver stages (50, 51). Overexpression of ATG8 in *Plasmodium berghei* merosomes leads to diffuse acyl carrier protein staining indicative of a collapse in apicoplast integrity (52). In *Toxoplasma gondii* which, like *P. falciparum*, is an apicomplexan parasite and important human pathogen, conditional knock-out of the apicoplast-localized two-pore channel protein (TgTPC) resulted in enlarged, vacuolated apicoplasts, reduced inter-organelle communication between the apicoplast and ER, and a significant decline in parasite fitness (53). A more recent investigation of *Toxoplasma* bradyzoites, the persister forms of *T. gondii* responsible for chronic infections, indicated that the apicoplast is vital for long-term survival of bradyzoites. Interestingly, tissue cysts derived from mutant parasites with a non-functional dynamin-related protein (Drp) display enlarged, reticulated apicoplasts and compromised persistence (54).

Parasite dormancy and recrudescence have been shown to occur irrespective of *pfk13* genotype, and *P. falciparum* parasites from repeated recrudescences after ART exposures, in vivo or in vitro, presented with no acquired *pfk13* mutations (33, 55). While recrudescent populations of the same response phenotype and genotype as the original population are characteristic of the dormancy phenomenon (56), it remains possible that dormancy may facilitate the acquisition of genetic mutations of drug resistance against ART or partner drugs of ACTs (57–59), analogously to the emergence of antibiotic resistance that has been described from persister cells in treated bacterial infections (60). In this regard, we note that selection of a PfK13 M476I mutation was reported from a Tanzanian isolate of *P. falciparum* after 30 cycles of ART pressure in vitro (61); however, no PfK13 mutations were obtained in two other long-term selection studies (62, 63) or in a separate study that identified mutations of the *pfcoronin* gene after 4 years of DHA pressure (64). In earlier studies of dormancy and parasite survival, Nakazawa *et al.* demonstrated recrudescences of *P. falciparum* populations treated with high levels of PYR, MEF, or sorbitol in vitro (9, 10) as well as recrudescences of *P. berghei* populations treated with CQ in vivo (15).

Dormancy and recrudescence of *P. falciparum* erythrocytic stages have also been reported after exposure to atovaquone alone or in combination with proguanil (17) and, more recently, to CQ in vitro (16). Taken together, these recrudescences of genetically distinct *P. falciparum* lines after treatment with different chemical agents, irrespective of the presence or absence of drug resistance mutations, are strong evidence for erythrocytic-stage dormancy as an innate survival mechanism.

ACTs require lasting action from effective partner drugs to prevent recrudescences from persisters after the powerful yet short-lived action of ART is done. First proposed in 1984, the use of a partner drug was introduced to control RI-type failure rates of up to 50% after monotherapy with ART alone (3–6). Variable parasite clearance times were also described in patients who received ART monotherapy (7), however careful studies found no associations between increased parasite clearance times and recrudescences or delayed resolutions of malaria fever after 2–7-day courses of monotherapy (7, 65–68)*. While the World Health Organization has defined ‘artemisinin partial resistance’ to include a parasite clearance half-life >5 hours after ART administration (72), there has been no confirmed evidence for progression from the original RI-phenotype to a RII or RIII level of clinical resistance (73) (see (58) for review). Several studies have highlighted the importance of an effective and durable partner drug to eliminate persisters after ART treatment. For example, in 2004, an International Artemisinin Study Group meta-analysis of CQ or amodiaquine in combination with AS (74) reported cure rates of 7–53% (28-day, PCR uncorrected) with the CQ-containing ACT in the Ivory Coast, Burkina Faso, and Sao Tome–Principe, whereas the PCR uncorrected rates for the amodiaquine-containing ACT were 68–85% in Kenya, Senegal, and Gabon. The combination of CQ with AS may have been compromised by a relative inability of CQ to kill the persisters (75), potential antagonism of CQ with AS (76), or emerging CQ resistance from the spread of PfCRT mutations in Africa (77). In southeast Asia, PPQ combined with DHA quickly failed due to resistance that resulted from more recent mutations in PfCRT (78–80). Similarly, antifolate mutations have compromised sulfadoxine-pyrimethamine (SP)-containing ACTs in regions of Africa, where alternative partner drugs such as lumefantrine or PPQ must now be used instead (81–84). Partner drugs able to efficiently prevent recrudescences from persisters remain critical, and treatment outcomes must be continually monitored for failures from mutant drug-resistant parasites.

Chloroplast swelling, vesiculation, and degradation with cell death are well-known responses to ROS and ROS-induced metabolites in a wide range of plant species (22). Could the various apicoplast morphologies we have described here reflect analogous responses that affect the survival of persisters? We note that ROS production is central to the action of ART, which uses an endoperoxide bridge to mediate parasite damage and death by generating free radicals that alkylate cellular components (85–88), and that ROS are involved in the toxicity of sorbitol to the cells of other systems (89–91). Considering the aberrant plastid morphologies that can occur in response to stressors in plants, *Toxoplasma*, and *Plasmodium* parasites (22, 45–49, 52, 53, 92–94), we speculate that swollen forms of the apicoplast may be associated with deterioration of the organelle and loss of persister viability. Sorting and characterization of persisters based on apicoplast morphology and mitochondrial activity paired with live-cell imaging and outgrowth experiments should help to uncover the fates of parasites with large and small apicoplasts. Elucidation of molecular mechanisms that govern organelle functions in dormancy may suggest new chemotherapeutic approaches to eliminate persisters and prevent recrudescences of malaria infections.

## Materials and Methods

### Parasite cultivation

*P. falciparum* NF54 parasites expressing a codon-optimized variant of green fluorescent protein, called Super Folder Green (SFG), fused to the N-terminus of the first 55 amino acids of Acyl Carrier Protein (ACP) were used to visualize the apicoplast (32). Parasite growth and maintenance was supported as previously described (28). Leukocyte-depleted human erythrocytes delivered weekly (Interstate Blood Bank, Memphis, TN) were washed three times with filtered incomplete RPMI 1640 (iRPMI, with 25 mM HEPES and 50 µg/ml hypoxanthine; KD Medical, Columbia, MD) and stored at 50% hematocrit at 4°C. Washed erythrocytes were used within a week of delivery and processing. Parasites were grown at 5% hematocrit, 37°C, and a gas mixture of 90% N2, 5% CO2, and 5% O2 in complete RPMI culture medium (cRPMI) composed of iRPMI 1640 with 1% AlbuMax I (Life Technologies, Carlsbad, CA), 0.21% sodium bicarbonate (KD Medical, Columbia, MD), and 20 µg/ml gentamicin (KD Medical, Columbia, MD). Parasitemia was monitored with Giemsa-stained thin blood smears and cultures were maintained below 5% parasitemia. Thin blood smears were fixed with 100% methanol (Sigma-Aldrich, St. Louis, MO) and stained for 15 minutes in 20% Giemsa (Sigma-Aldrich, St. Louis, MO; diluted in deionized water).

### Dihydroartemisinin treatment and osmotic removal of mature parasite stages

To obtain synchronized ring-stage parasites prior to DHA exposure, cultures were treated twice for 10 min with 10 ml of 5% D-sorbitol (Thermo Fisher Scientific, Ward Hill, MA), 46 h apart. Each treatment occurred at room temperature, lasted 10 minutes, and was followed by one wash with iRPMI (95). Directly after the second sorbitol treatment, cultures were diluted to 2% parasitemia and 5% hematocrit at a total volume of 10 ml. A stock solution of 700 µM DHA (Sigma-Aldrich, St. Louis, MO) dissolved in DMSO (Sigma-Aldrich, St. Louis, MO) was diluted in cRPMI to treat parasites at a final concentration of 700 nM DHA/0.1% DMSO for 6 h at 37°C. In parallel, a separate culture was treated with 0.1% DMSO diluted in culture medium as a control. Cells of both treatments were washed twice with cRPMI and returned to culture conditions. Blood smears were prepared prior to drug treatment and following the final wash. 24 h post-DHA, cells were subjected to 5% sorbitol for 30 minutes at 37°C to eliminate mature parasites that survived DHA treatment. Simultaneously, cells that were exposed only to DMSO underwent a mock-sorbitol treatment with iRPMI. Following two washes with iRPMI, cells were returned to cRPMI under normal culture conditions.

### Dihydroartemisinin and sorbitol treatment and monitoring of cultured parasites for recrudescence

Parasite cultures were synchronized and treated with the vehicle control or DHA as described above. At 24, 48, and 72 h post-drug pulse, both DHA and vehicle-treated cultures were subjected to 10 ml of 5% sorbitol for 30 minutes at 37°C, followed by two washes with iRPMI. The cells were returned to culture in new T25 flasks with complete medium following each treatment. After the last sorbitol treatment at 72 h, 200 µl of 50% hematocrit stock red blood cells were added to replenish those lost due to lysis. Media was changed daily, and 100 µl of 50% hematocrit stock red blood cells were added every third day. Recrudescence was monitored via daily thin blood smears until 2% parasitemia was reached.

### Fluorescence-activated cell sorting

Two days (t= 50 h) after treatment with DHA/sorbitol or the vehicle control, 1 ml of culture was removed from each flask and centrifuged at 3,000 rpm. Pelleted cells were washed three times with wash buffer composed of filter-sterilized 1× Hanks Balanced Salt Solution (HBSS; Gibco, Ward Hill, MA) and 2% fetal bovine serum (Gibco, Ward Hill, MA). Parasite DNA and mitochondria were stained using 2 µM Hoechst 33342 and 0.1 µM MitoTracker Deep Red FM (Invitrogen, Waltham, MA). Staining, which occurred at 37°C in the dark for 30 minutes, was followed by three wash steps in which cells were centrifuged and resuspended in 2 ml of wash buffer to remove excess stain.

Cells were sorted using the BD FACSAria Fusion fitted with a 70-micron nozzle, and gating was performed using the BD FACSDiva software (BD Biosciences, Franklin Lakes, NJ) Samples underwent two sequential sorting steps, herein called the yield sort and purity sort. In the yield sort, Hoechst fluorescence was used to separate parasitized red blood cells from uninfected red blood cells. Infected red blood cells collected in the yield sort were further enriched based on increasing MitoTracker fluorescence intensity in the purity sort to distinguish between erythrocytes harboring persisters or active parasites. Persisters and active ring-stage parasites were collected into 5 ml polypropylene round-bottom tubes (Falcon) using a 4-way sorting method.

Gating strategies were validated by using collected cells to make cytospins. 50,000-150,000 cells from each gate were diluted in 100-150 µl of FACS wash buffer and added to Epredia cytofunnels (Fischer Scientific, Catalog No. 59-910-40, USA) with mounted microscope slides. The cytofunnels plus mounted microscope slides were centrifuged in the Cytospin™ 4 Cytocentrifuge (Epredia™) for five minutes at 500 rpm. Funnels were removed and discarded, and the microscope slides were allowed to air-dry. Slides were then fixed in 100% methanol and stained in freshly prepared 20% Giemsa for 15 minutes.

### Airyscan microscopy analysis

Cells collected from FACS were transferred to a 1.5 ml tube (Eppendorf) and centrifuged. The pellet was resuspended in 30-300 µl of residual wash buffer depending on the number of cells collected from FACS. Clear silicon gaskets (Grace Bio-Labs, Bend, Oregon) were placed in the wells of Lab-Tek chambered cover glass with No. 1 borosilicate glass bottoms (Thermo Fisher Scientific, Ward Hill, MA) to create microwells 3 mm in diameter. 10 µl of 0.01% poly-L-lysine solution (Sigma-Aldrich, St. Louis, MO) was added to each microwell, incubated at room temperature for 30 minutes, and washed twice with 10 µl of 1× HBSS. Immediately after the second wash, 10-30 µl of the recovered cells were added to the poly-lysine coated microwells and incubated at room temperature for 30 minutes in the dark. Approximately 1.5 ml of a 5-fold dilution of Intracellular Fixation Buffer (Invitrogen, Waltham, MA; 1 part diluted into 4 parts by volume of 1× phosphate buffered saline pH 7.4) was added to each chamber, such that it covered the bottom of the chamber and microwells. Parafilm was used to cover the chambered cover glass before being stored at 4°C in the dark.

Within 24 h of fixation, parasites and their organelles were imaged using the Zeiss 880 LSM with Airyscan equipped with a Zeiss plan apochromat 63x/1.4 NA oil immersion objective. Individual organelles were segmented from Airyscan-processed z-stacks covering the entire organelle volume using threshold values determined via the Otsu method in FIJI/ImageJ. Segmented images were then used to construct three-dimensional surfaces in Imaris 9.7.2. Where there was ambiguous background signal, it was subtracted from the channel of interest to improve Otsu thresholding. Organelle volumes were obtained from constructed three-dimensional surfaces, and inter-organelle distances were calculated as the distance between the centers of each surface.

## Acknowledgments

We thank Jianbing Mu, Daniel Kiboi, Johannes S. P. Doehl, and Akshaykumar Nayak for discussions. This work was supported by the NIAID Division of Intramural Research.

## Data Availability Statement

The 717 Airyscan-processed images collected and analyzed in this study are publicly available at Bioimages Archive repository (https://www.ebi.ac.uk/biostudies/bioimages/studies/S-BIAD987). Organelle volume information and distance values with summary statistics are provided in supplementary **Dataset S1** accompanying this report.

---

**Figure S1.**
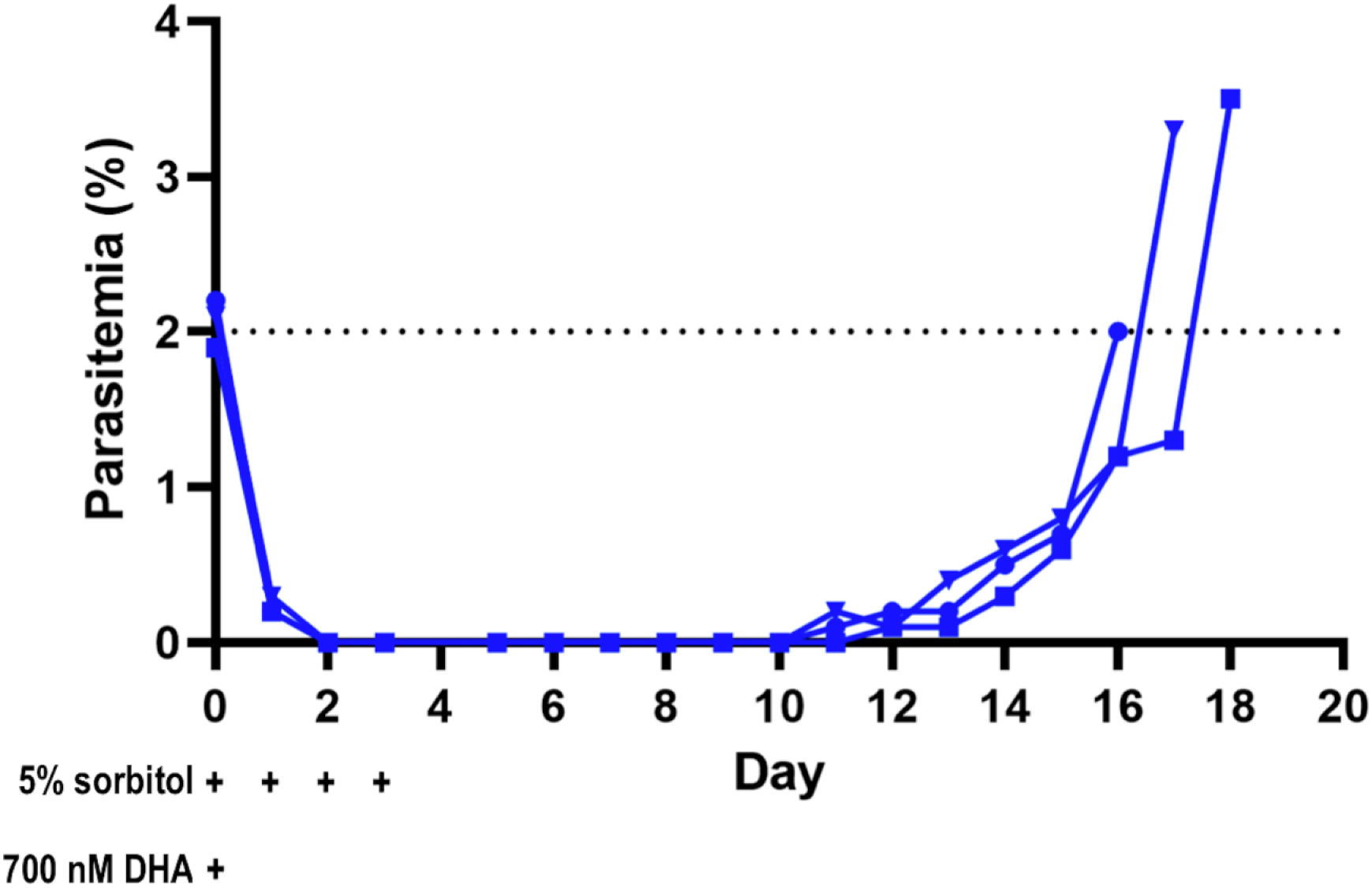
Recrudescence curves of *P. falciparum* NF54-SFG active-form blood stages treated with 700nM DHA and sorbitol selection in vitro. Highly synchronized NF54-SFG ring-stage parasites carrying Super Folder Green (SFG) fused to the leader sequence of the acyl carrier protein (ACP) were treated with 700 nM DHA for 6 h. At 24, 48, and 72 hours after DHA treatment, the parasites were subjected to selection with 5% sorbitol for 30 minutes (indicated by +). Parasitemia of active forms including ring-, trophozoite and schizont-stage parasites was monitored daily by Giemsa-stained thin blood smears until ≥2% recrudescence. Each blue line represents one biological replicate (n=3).

**Figure S2.**
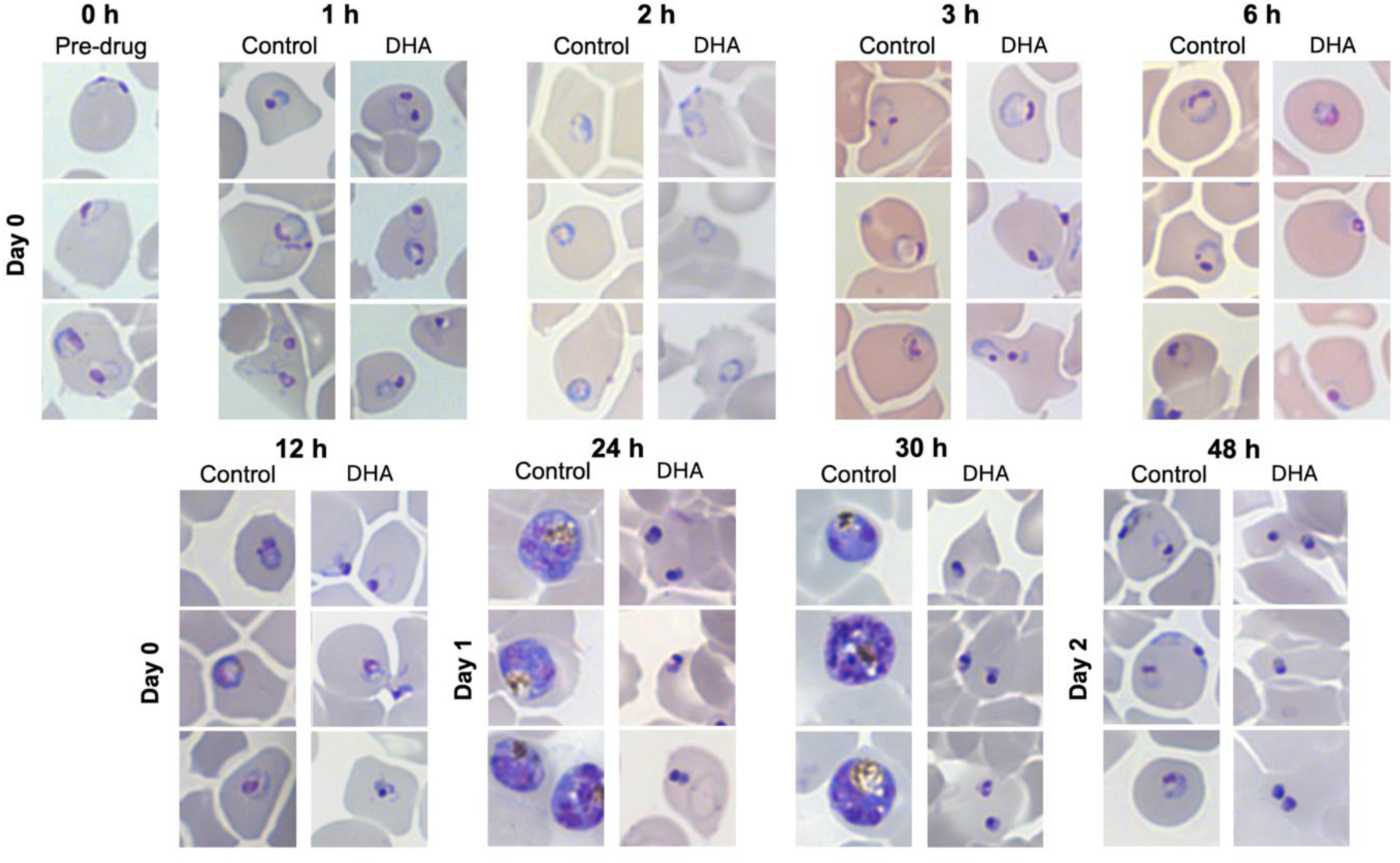
Images of Giemsa-stained NF54-SFG parasites before, during, and after drug treatment. Synchronized cultures consisting of 6-9 h ring-stage parasites (0 h) were treated with 700 nM DHA or the vehicle control for 6 h on day 0. Ring forms were predominant at 1, 2, 3 and 6 h in both the control- and DHA-treated conditions. At 12 h, early trophozoites were present in control-treated cultures while DHA-treated parasites displayed abnormal morphology. On day 1 at 24 h and 30 h, persister forms were apparent in DHA-treated cultures. Control-treated cultures consisted mostly of late trophozoites, schizonts, and very early rings at 24-30 h. By 48 h, the control-treated population was comprised mostly of rings and the DHA-treated population contained persisters. Three representative images are shown for each timepoint and treatment.

**Figure S3.**
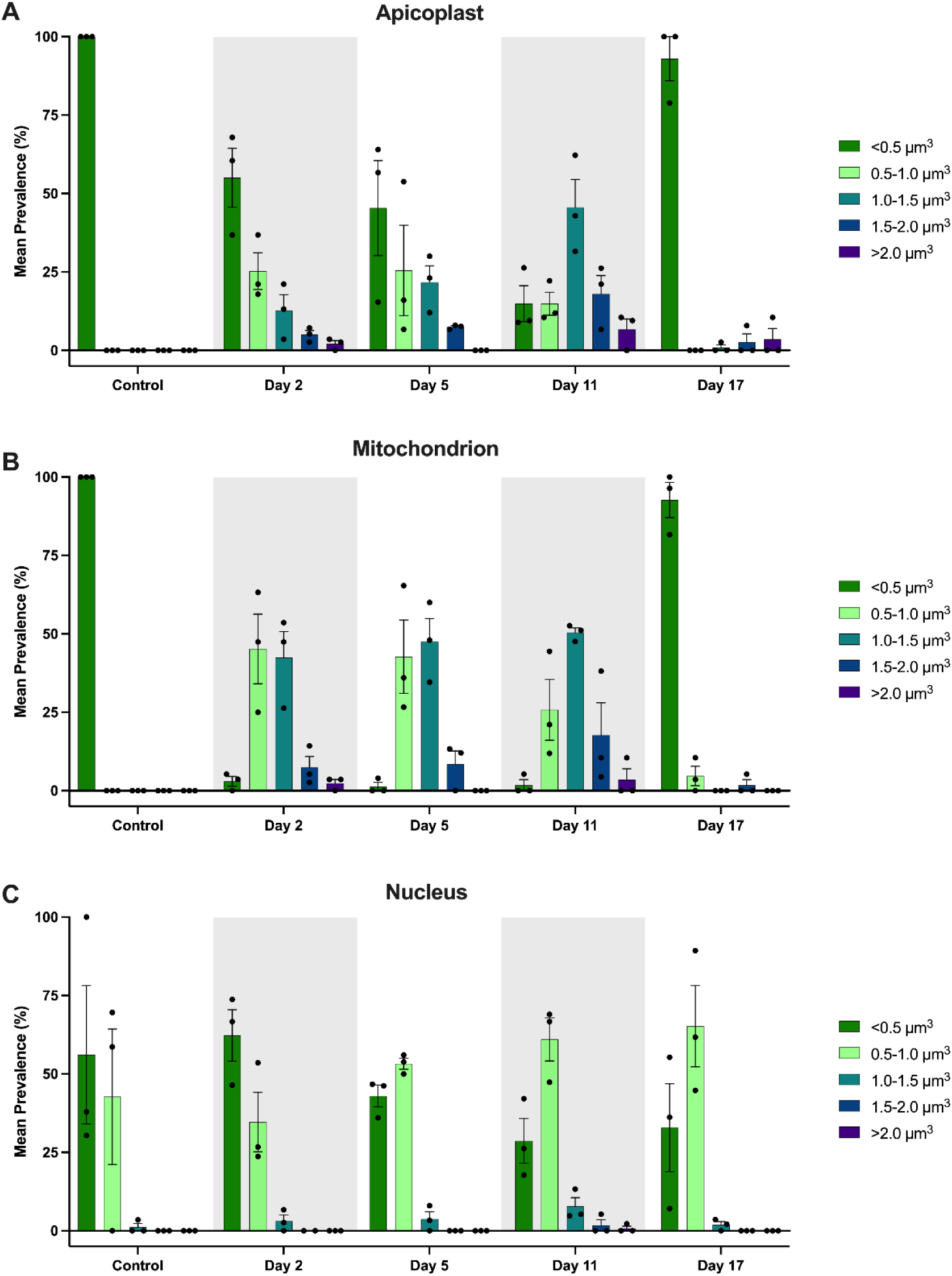
Distribution of organelle volumes following DHA/sorbitol treatment. Histograms display the mean prevalence and standard error of mean (SEM) of apicoplast **(A)**, mitochondrion **(B)**, and nucleus **(C)** volumes in control-treated parasites and DHA/sorbitol-treated parasites on days 2, 5, 11, and 17 obtained from three biological replicates. Organelles were sorted into five groups based on volume: <0.5 µm^3^, 0.5-1.0 µm^3^, 1.0-1.5 µm^3^, 1.5-2.0 µm^3^, and >2.0 µm^3^.

**Movie S1 (separate file).** 3D rotated view of well-separated organelle fluorescence and surfaces of a vehicle control-treated ring-stage parasite. The movie first displays apicoplast (green), mitochondrion (red), and nucleus (blue) fluorescence volumes assembled from Airyscan-processed z-slices. Fluorescence volumes are then presented concomitantly with transparent Imaris surface renderings. Finally, only the transparent surface renderings are shown. The organelles presented here correspond to the first image of the second row under “Vehicle control” in Figure 3A. Scale bar indicated in movie.

**Movie S2 (separate file).** 3D rotated view of organelle fluorescence and surfaces in close approximation in a vehicle control-treated ring-stage parasite. The organelles presented here correspond to the second image of the second row under “Vehicle control” in Figure 3A. Scale bar indicated in movie.

**Movie S3 (separate file).** 3D rotated view of organelle fluorescence and surfaces of a DHA/sorbitol-treated persister with a regularly sized, oblate apicoplast. The organelles presented here correspond to the first image of the second row under “DHA/sorbitol” in Figure 3A. Scale bar as indicated in movie.

**Movie S4 (separate file).** 3D rotated view of organelle fluorescence and surfaces of a DHA/sorbitol-treated persister with a large and irregularly shaped apicoplast. The organelles presented here correspond to the second image of the second row under “DHA/sorbitol” in Figure 3A. Scale bar as indicated in movie.

**Dataset S1 (separate file).** Workbook with all quantitative data presented in this manuscript. Organelle volumes and distances represented in Figure 2B and C, Figure 3B and C, Figure 4, and Figure 5C and D are included. Results of statistical analysis for these figures are also included.

## Footnotes

* More recent reports have described regional associations *pfk13* mutations with longer clearance times after ART treatment (69–71), but these reports do not address or challenge the findings of references (7, 65–68), nor do they provide evidence for *pfk13* marker associations with greater recrudescence frequencies or prolonged fever that would indicate increased clinical resistance to the ART component of ACT.

